# Unraveling the effects of polyhydroxyalkanoates accumulation in *Pseudomonas extremaustralis* growth and survival under different pH conditions

**DOI:** 10.1101/2024.06.19.599761

**Authors:** María Gabriela Brito, Nancy I. López, Laura J. Raiger Iustman

## Abstract

Polyhydroxyalkanoates (PHAs) are intracellular polymers that enhance bacterial fitness against various environmental stressors. *Pseudomonas extremaustralis* 14-3b is an Antarctic bacterium capable of accumulating, short-chain-length PHAs (sclPHAs), composed of C3–C5 monomers, as well as medium-chain-length PHAs (mclPHAs) containing ≥C6 monomers. Since pH changes are pivotal in bacterial physiology, influencing microbial growth and metabolic processes, we propose that accumulated PHA increases *P. extremaustralis* fitness to cope with pH changes. To test this, we analyzed the production of sclPHA and mclPHA at different pH levels and its effect on bacterial survival against pH stress. *P. extremaustralis* was able to grow and accumulate PHA when the culture media pH ranged from 6.0 to 9.5, showing a marked loss of viability outside this range. Additionally, based on the analysis of different PHA-deficient mutants, we found that when exposed to both acidic and alkaline conditions, sclPHA and mclPHA conferred different protection against pH stress, with sclPHA making the main contribution. These results highlight the importance of PHA in supporting survival in pH-stressful environments.

## 1 Introduction

Bacteria are constantly exposed to fluctuating environmental conditions such as temperature, oxygen, nutrient availability, and pH. Specifically, pH was described to affect microbial growth by influencing the chemical activity of protons, having a key role in redox reactions, and interfering with bacterial metabolism [1]. Several mechanisms were proposed to explain how bacteria can sense and cope with pH changes in the environment, most of them related to changes in the outer membrane, proton efflux, and ATPase activity [2]. In addition to these specific mechanisms, other direct or indirect features may help bacteria cope with stress arising from pH fluctuations short-chain-length PHAs (sclPHAs), composed of C3–C5 monomers, as well as medium-chain-length PHAs (mclPHAs) containing ≥C6 monomers *Pseudomonas* species. Nevertheless, some *Pseudomonas* species belonging to *P. oleovorans* group’ and others like *Pseudomonas* sp. 61-3, *P. extremaustralis* 14-3b, and *P. pseudoalcaligenes* can accumulate both scl and mclPHAs [8-11]

It is known that bacterial growth is influenced by various factors such as temperature, carbon, nitrogen, phosphorus, and pH. However, the study of the influence of pH variation on PHAs production has been limited mostly to optimizing culture conditions in different producing bacteria, mainly belonging to *Bacillus* genus, recombinant *E. coli* or producers consortia [12-18]. Moreover, the influence of PHA in pH-induced stress conditions has been even less explored [19].

*P. extremaustralis* is an extremophile bacterium isolated from a temporary pond in Antarctica [20], showing high-stress resistance associated with PHAs production [21, 22]. This bacterium can produce and accumulate mostly sclPHAs, but also small amounts of mclPHAs [23]. For this reason, *P. extremaustralis* serves as a good model to study how environmental pH influences the production of the different kinds of PHAs, and the role of these polymers in coping with pH stress. This work aims to analyze the effect of PHA accumulation on bacterial growth and survival in response to pH challenges and the contribution of different kinds of PHA to the survival of *P. extremaustralis* under pH changes.

## 2 Materials and Methods

### 2.1 Strains and culture media

*Pseudomonas extremaustralis* 14-3b DSM 25547 and their polyhydroxyalkanoate deficient mutants PHB-, PHA- and PHAB-(Table 1) were cultured in Lysogeny broth medium (LB) supplemented with 0.25% sodium octanoate (SIGMA) (LBO) which lead to PHA accumulation. When necessary, kanamycin (Km), gentamicin (Gm), or tetracycline (Tet) was added at a concentration of 20 µg/ml, 10 µg/ml, and 10 µg/ml respectively. For the construction of the double mutant of *P. extremaustralis* PHAB-we used the PHA-mutant (*ΔphaC1ZC2*::Gm) previously built in the laboratory [24] and the *ΔphbC*::Km fragment that comes from the PHB^-^ mutant inserted in the suicide vector pPHU281, encoding tetracycline-resistance [25]. The recombinant plasmid was transferred by electroporation to *E. coli* S17-1. Then, the conjugation of *P. extremaustralis* PHA-mutant and *E. coli* S17‐1 harboring the pPHU281*ΔphbC*::Km was carried out. For the selection of transconjugant strains, we used 0.5NE2 medium [26] supplemented with Gm, Km, and sodium octanoate as a carbon source where the donor bacteria cannot grow. Double homologous recombination resulted in tetracycline sensible clones with the *phbC*::Km inserted at the target locus in the chromosome. The mutant was also verified by sequencing using primers external to the construction (data not shown). Table 1 resumes the relevant genotype and phenotype of the strains used in this work.

**Table 1.** Characteristics of the bacterial strainsTable 1. Characteristics of the bacterial strains.

| Strain | Relevant genotype/ phenotype | Reference |
| --- | --- | --- |
| *P. extremaustralis* 14-3b<br>DSM 25547 | WT, sclPHA and mclPHA<br>producer | [10, 20] |
| PHB- | $\Delta phbC::Km$ | [22] |
| PHA- | $\Delta phaC1ZC2::Gm$ | [24] |
| PHAB- | $\Delta phaC1ZC2::Gm$, $\Delta phbC::Km$ | This work |

### 2.2 Bacterial growth conditions

For all experiments, a -80°C glycerol conserved stock of each strain was plated in LB and cultured at 28°C for 24h. Pre-cultures or “starter cultures” were prepared using isolated single colonies for each strain, which were used to inoculate 5 ml of LBO in 50 ml bottles and incubated at 28°C and 180 rpm overnight. Independent pre-cultures were used to inoculate cultures with different initial pH. For these cultures, the LB medium was adjusted to the following pH: 5.5; 6.0; 7.0; 7.5; 8.0; 9.0; 9.5; 10.0, or 11.0 by addition of either 1 M HCl or 1 M NaOH as necessary, before sterilization by filtration. After filtration, sterile sodium octanoate was added and pH was verified.

### 2.3 Growth at different pH

For the assays, 100 ml Erlenmeyer flasks containing 10 ml of pH-adjusted LBO were inoculated with the starter cultures described above at an initial OD_600nm_=0.05. Immediately after inoculation, to achieve the effect of pH at the initial time (T0), a sample of 0.1 ml was collected to determine the number of viable bacteria, as colony-forming units (CFU)/ ml, in LB plates. The cultures were incubated at 28°C and 180 rpm for 24h. After incubation, the final OD and the CFU/ml were measured. Finally, 10 ml of each culture was centrifuged at 7000 rpm. The pH of the cell-free supernatant was measured and the cell pellet was stored at -80°C until gas chromatograph (GC) analysis.

### 2.4 PHAs production

PHA production was analyzed both qualitatively and quantitatively. Qualitative analysis was carried out using Nile Blue A staining. In this case, samples of each culture were stained with 1% Nile blue [27] and observed in an epifluorescence microscope at an excitation wavelength of 460 nm.

Quantitative analysis of PHAs production and monomer composition was determined as previously described [28]. Briefly, lyophilized cell pellets were subjected to methanolysis [29].

Monomers methyl esters were analyzed using an Agilent 7820 gas chromatograph (GC) equipped with a flame ionization detector (FID), automatic liquid sampler ALS 7693, and an HP-5 capillary column. Standard solutions were prepared using poly (3-hydroxybutyric acid) (SIGMA Aldrich) and mcl-PHAs from *Pseudomonas putida* KT2440. Benzoic acid (0.5 mg/ml) was used as an internal standard. Triplicate experiments from independent cultures were performed with three GC measurements for each sample. PHAs content was calculated as the percentage of cell dry weight (CDW). The total content of sclPHA and mclPHA was determined by adding all the detected short-chain and medium-chain monomers. For sclPHA, this comprised almost exclusively PHB, while in the case of mclPHA, C6 and C8 monomers were detected, with C8 being the most abundant.

### 2.5 pH Tolerance

The effect of PHAs accumulation in the resistance to different pH was analyzed using a disc inhibition assay as previously described [21]. Briefly, 0.1 ml of 24h LBO cultures (pH=7.5) were spread into LB agar plates (20 ml LB agar) and a drop of 10 μl of different pH buffers was added on Whatman filter discs (6 mm diameter), placed immediately after inoculation, in the center of the plate. This assay was made by triplicate using independent cultures. The desired pH was achieved by using different buffer solutions. To pH 5.5: 0.1 N Sodium Acetate-Acetic acid buffer; pH 6.0 to 9.5: 0.1 N phosphate buffer and pH 10: 0.2 M Glycine-NaOH buffer. All the buffers were sterilized by filtration before use. LB agar plates were cultured at 28°C for 24-48h and the inhibition halo was measured.

### 2.6 Transmission electron microscopy (TEM)

The cell shape and the PHA granules of 24h LBO cultures of *P. extremaustralis* and its PHB^-^, PHA^-^ and PHAB^-^ mutants were analyzed by transmission electron microscopy (TEM). Cells were harvested, washed twice in PBS, and fixed in 2.5% (w/v) glutaraldehyde in the same solution. Then, cells were suspended in 2.5% (w/v) OsO_4_ for 1 h, gradually dehydrated in ethanol (50%,70%, 96%, and 100% (v/v) for 30 min each, completed by acetone for 20 min. Cells were embedded in “Durcupan” epoxy resin. Ultrathin sections (thickness 70-90 nm) were made with an Ultramicrotome (Reichert Jung Ultracut E) using a glass knife. Finally, the sections were mounted on copper grids, contrasted with uranyl acetate and lead citrate (method of Reynolds), and examined in a Zeiss EM109 T. Digital photographs were taken with a Gatan ES 1000 W digital camera. Analyses were performed using LANAIS-MIE Institut service (Faculty of Medicine, UBA-CONICET).

### 2.7 Statistical analysis

The significance of each treatment was evaluated by Student’s t-test with confidence levels at 95% (i.e., P < 0.05 was considered as significant).

## 3 Results

### 3.1 The effect of different pH on *P. extremaustralis* growth

LBO cultures adjusted to pH 5.5 to 11.0 were inoculated with the pre-cultures. The number of viable cells (CFU/ml) and OD_600_ were measured immediately post-inoculation (T0) and after 24 h of culture (T24) (Fig. 1). All cultures started at an OD_600_ 0.05 corresponding to approximately 1x10^7^ CFU/ml. When the inoculum was added into LBO at pH 5.5 or pH 11.0 (T0), a marked viability decrease, of about 5 orders of magnitude, was detected. This loss of viability was not observed for cultures adjusted at pH 6.0, 7.0, 8.0, 9.0, 9.5, and 10.0 (Fig. 1). In line with this, after 24 hours, cultures developed in LBO initially adjusted to pH levels between 7.0 and 9.5 showed an average growth increase of about 3 orders of magnitude (Fig. 1). Acidic conditions, at pH 5.5 and 6.0, negatively impacted cellular growth, with the culture at pH 5.5 being the most affected (Fig. 1). Although cells inoculated at pH 6.0 showed no effect at T0, final growth was indeed affected, reaching only 3.5 10^8^ CFU/ml at 24 h (Fig. 1). Similar results were achieved at alkaline conditions (pH 10.0). At pH 11.0 no counts were observed at T24 (Fig. 1). Similar results to CFU counts were observed regarding OD_600_ measurements (data not shown).

**Fig. 1.**
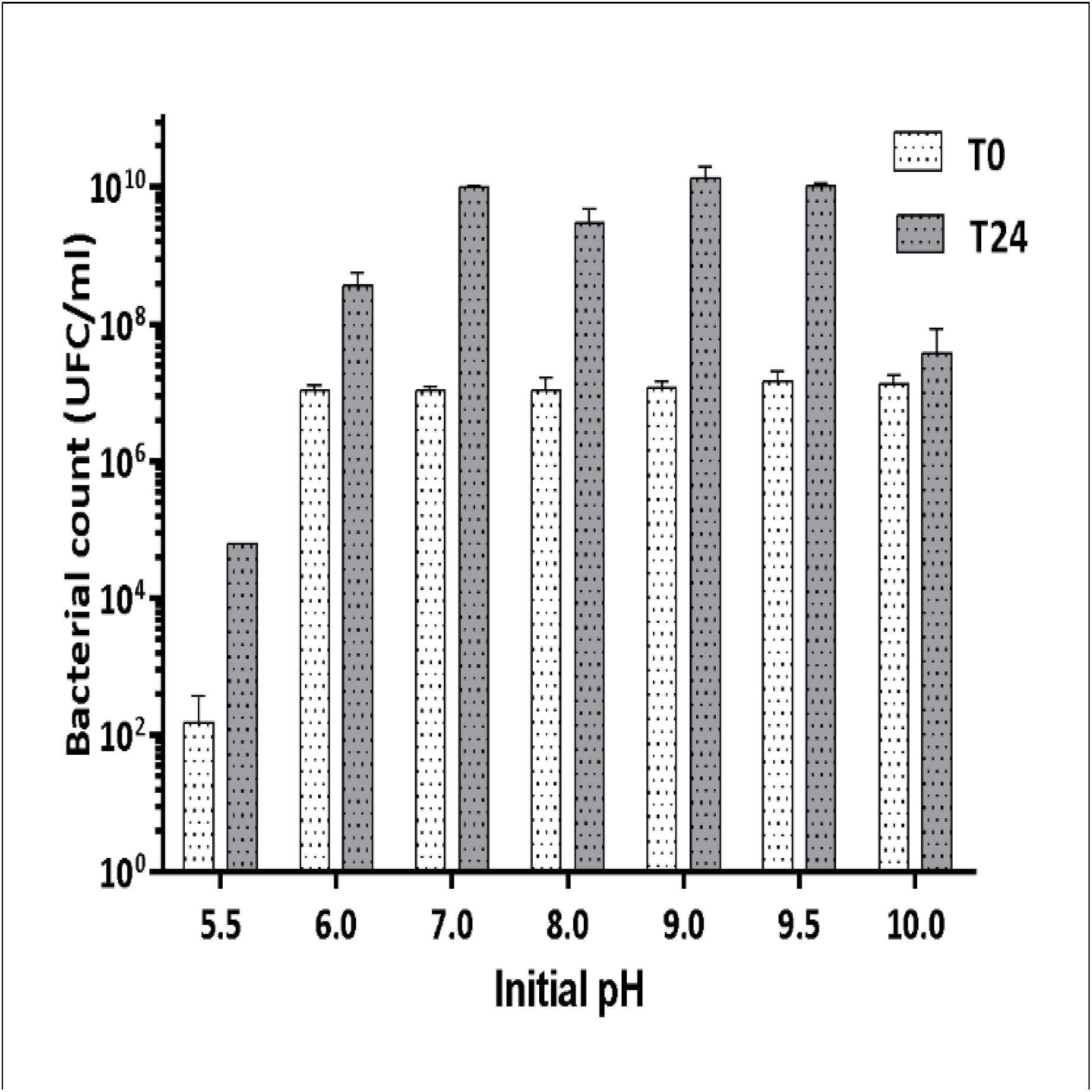
Effect of different pH on *P. extremaustralis* growth. The viable bacterial number at the initial time (T0) and after 24h culture (T24). Values represent the mean ± SD of triplicate independent cultures.

The pH of the cell-free supernatant at the end of the experiments was also measured. Cultures whose initial pH ranged from 7.0 to 9.5 reached a final pH of around 8.6. On the other hand, the cultures inoculated at an initial pH of 5.5, showed almost no changes at T24. Finally, those grown at pH 6.0 increased the culture medium pH to 7.0, while the cultures grown at pH 10.0 showed a slight pH decrease (Fig. 2).

**Fig. 2.**
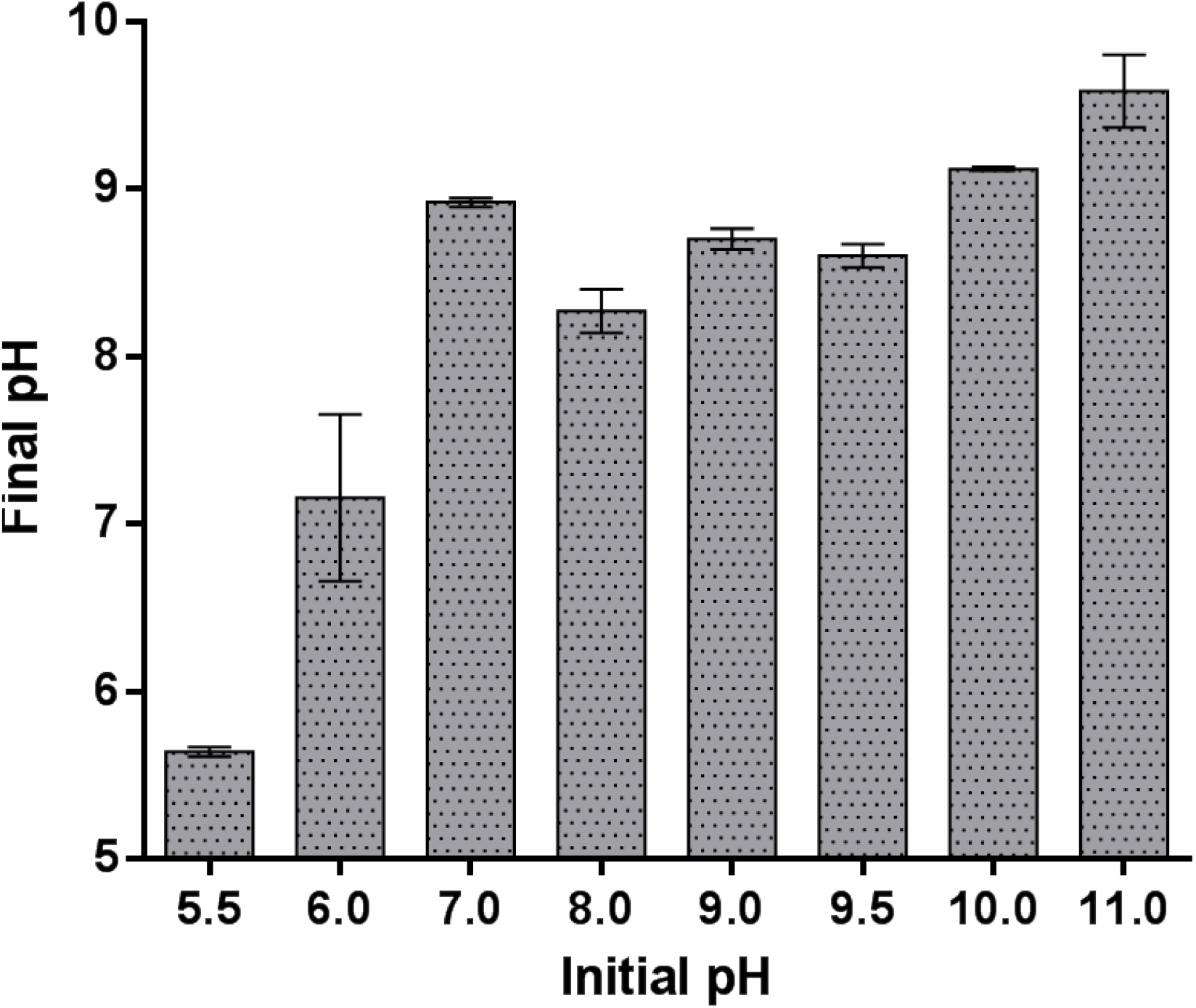
pH of cell-free supernatants of *P. extremaustralis* cultures developed with different initial pH for 24h. Values represent the mean ± SD of triplicate independent cultures

TEM analysis of the starter cultures showed notable heterogeneity in the PHA accumulation pattern (Fig. 3A). Several cells presented large PHA granules, while others had small granules or none at all (Fig. 3B). This pattern was observed in all the replicates. Nevertheless, only small variability among replicates was observed when total PHA content was measured (36.72 ± 1.24% CDW).

**Fig. 3.**
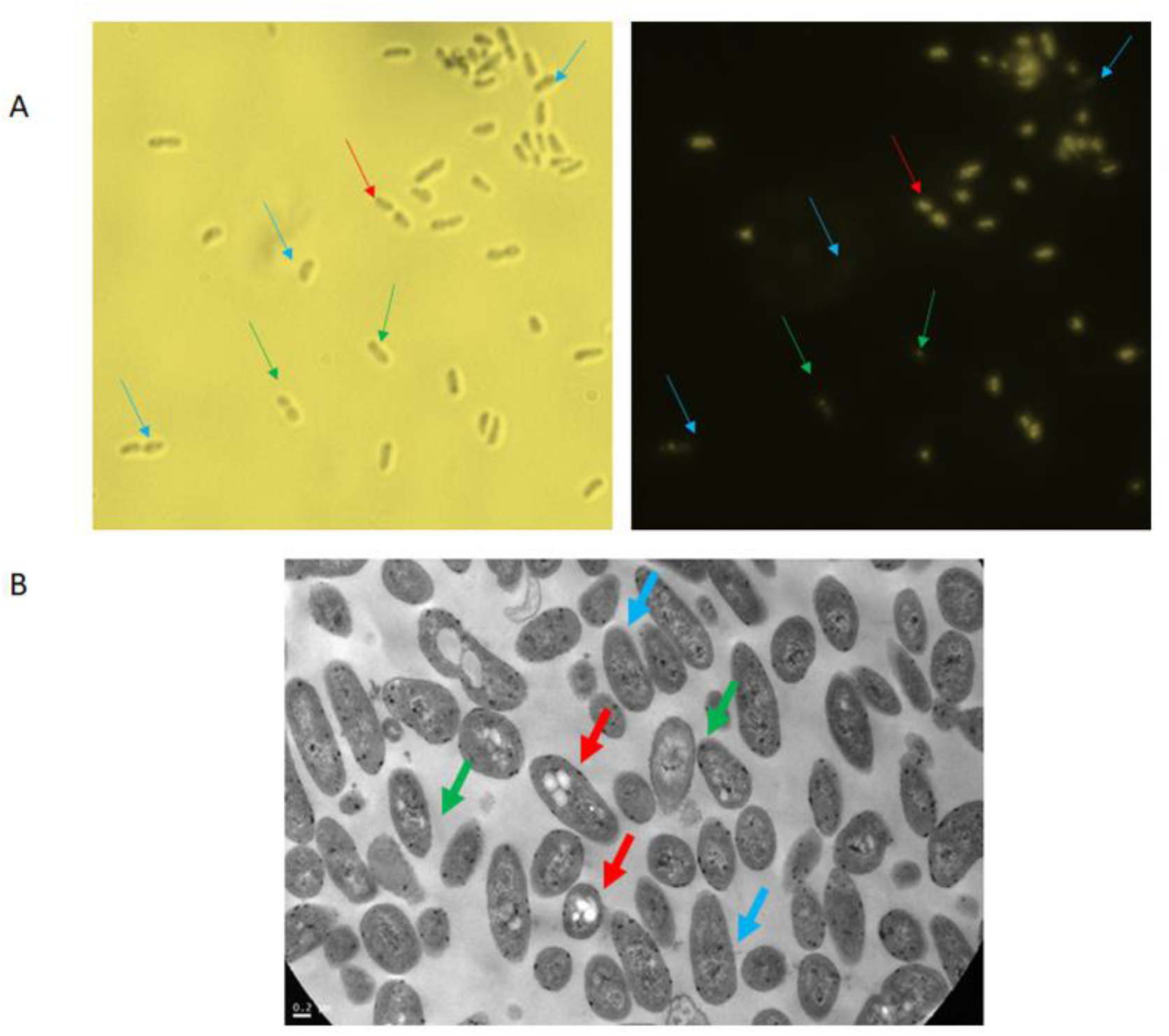
PHA accumulation of cells in starter cultures. A) Microscopic image of Nile blue stained cells after illumination with white light (left) or blue light, 460 nm (right). Magnification 1000X. B) TEM images showing different PHA content. Arrows indicate PHA accumulation range: high (red), intermediate (green), or low (light blue).

### 3.2 Effect of pH on PHA Accumulation

*P. extremaustralis* was able to accumulate PHAs when the initial pH was adjusted to 6.0, 7.0, 7.5, 8.0, 9.0, and 9.5, with cells showing rod shape and good development (Fig. 4A). When it was grown at pH 5.5 and 10.0, a scarce number of small round-shape cells was observed, showing evidence of stress. None of them showed PHA granules.

**Fig. 4.**
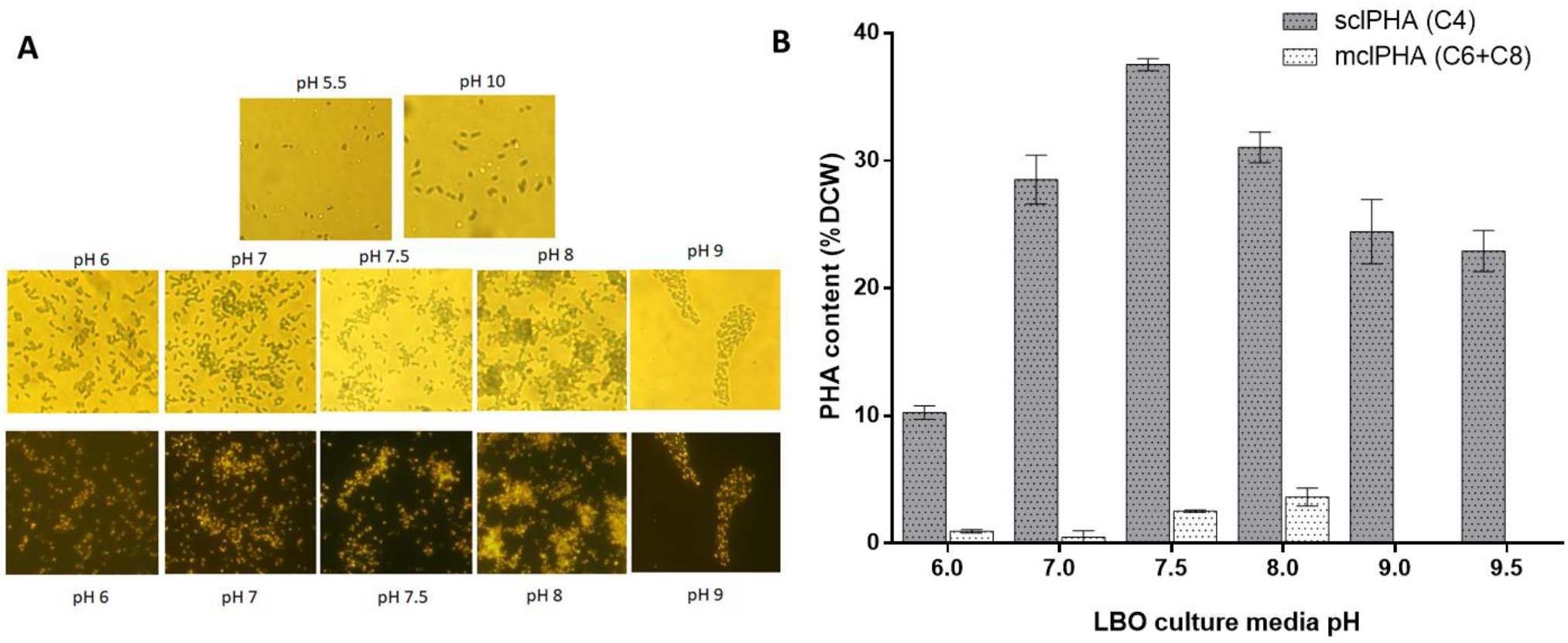
**A) Microscopic observation** of Nile Blue A stained *P. extremaustralis* cells after 24h of culture in LBO medium initially adjusted at different pH. Top: bright field microscopy (1000x) and bottom: epifluorescence microscopy (1000x). B). **PHAs production** of P. extremaustralis grown at different initial pH in the culture medium. sclPHA: short chain length PHA; mclPHA: medium chain length PHA Bars represent the mean + SD of triplicate independent cultures. CDW: Cellular dry weight.

GC analysis showed that at an initial pH of 7.5, *P. extremaustralis* was capable of producing both mclPHA and sclPHA, reaching the highest sclPHA accumulation (37.54 ± 0.47 % C DW, Fig. 4B). The PHB content was significantly lower at pH 6.0 than those achieved at pH values ranging from 7.0 to 9.5 (t-test, P<0.05, Fig. 4B). At more alkaline pH values (9.0 and 9.5) a significant decrease in sclPHA content was observed in comparison with polymer content obtained at pH 7.0 to 8.0 (P<0.05). On the other hand, mclPHAs were detected at pH 6.0, 7.0, 7.5, and 8.0, with cells grown at pH 6.0 and pH 8.0 showing the lowest and highest amount, respectively (P<0.05, Fig. 4B).

### 3.3 Tolerance to pH stress

To determine if the accumulation of polymers could be responsible for the bacterial fitness against acid and alkaline stress, the wild-type strain (WT) and three PHA-deficient mutants were exposed to different pH in a disc inhibition test. Representative TEM microphotographs of PHA granules in the WT and mutant strains can be observed in Fig. 5A.

**Fig. 5.**
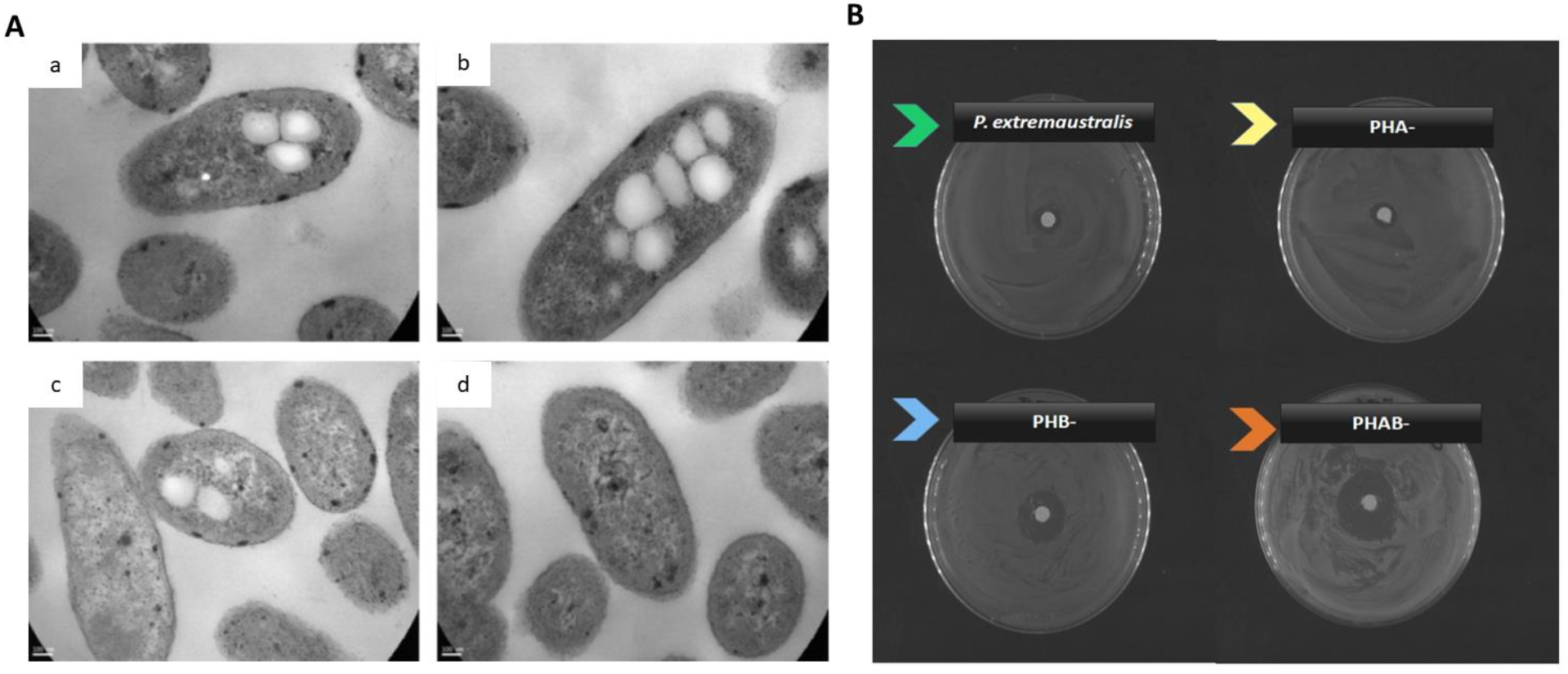
**A)** Transmission electron microscope (TEM) observations of PHA granules in *P. extremaustralis* and mutant strains. a: WT, b: PHA-, c: PHB- and b: PHAB-strains. Bar=100 nm, 85000 X. **B)** A representative photograph of one of the replicates showing the sensitivity to acidic (5.5) pH of *P. extremaustralis* and PHA deficient mutants assessed by disc diffusion test.

These analyses involving the use of different mutant strains allowed the discrimination of the effect of sclPHA and mclPHA, and those due to the lack of PHA under the same physiological conditions. Table 2 shows the PHA accumulation in each strain and the response to pH 5.5 and 10.0 buffer solutions, where the WT strain resulted in more sensitivity to alkaline pH than to acidic pH (P<0.05). No inhibition halo was observed around the disc corresponding to the buffer solution adjusted to pH 6.0 to 9.5 (data not shown). At pH 5.5 the strain unable to synthesize sclPHA showed a larger inhibition halo than the WT (P<0.05). The acidic pH effect was strongest in the PHAB^-^ mutant, unable to produce any kind of polymer in comparison with the WT (P<0.05). Representative images corresponding to the inhibition test at pH 5.5 of *P. extremaustralis* and its mutants are shown in Fig. 5B. At pH 10.0, the three mutants showed a larger inhibition halo than the WT (P<0.05, Tabla 2).

**Table 2.** PHA production in *P. extremaustralis* and mutant strain used for the pH tolerance test and inhibition halo after exposure to acidic or alkaline pH.

| Strain | PHA composition (% CDW) |  | Inhibition halo (cm) |  |
| --- | --- | --- | --- | --- |
|  | C4 | C6+C8 | pH 5.5 | pH 10 |
| WT | 36.72 ± 0.95 | 2.60 ± 0.06 | 1.04 ± 0.08 | 1.30 ± 0.02 |
| PHA- | 22.81 ± 0.52 | Not produced | 1.09 ± 0.06 | 1.51 ± 0.09 |
| PHB- | Not produced | 3.68 ± 0.06 | 1.66 ± 0.25 | 1.77 ± 0.05 |
| PHAB- | Not produced | Not produced | 2.62 ± 0.12 | 2.03 ± 0.13 |

## 4 Discussion

Several studies have analyzed the PHA metabolism in different culture conditions, and their role in survival and resistance to several stress factors [4]. Nevertheless, to our knowledge, the effect of pH on *Pseudomonas* growth and its relationship with PHA metabolism have been less studied. *P. extremaustralis* was isolated from a temporary pond in Antarctica, arousing interest because of its ability to produce different types of PHAs and its fitness to cope with different stresses. In this work, we demonstrated that *P. extremaustralis* was able to develop within a range of pH from 6.0 to 9.5, showing PHA accumulation. Growth in this pH range was also observed in different *Pseudomonas* strains [30, 31], like *P. toyotomiensis*, a psychrotolerant species isolated from a hot spring, which can grow in a pH range from 6.0 to 10.5 [32]. An interesting observation was that *P. extremaustralis* viability immediately decreased when the pH of the culture media was adjusted to 5.5 or 11.0. Despite the loss of viability, slight culture growth was observed at pH 5.5 after 24 h, but no growth was obtained in cultures at pH 11.0. The persistence of *Pseudomonas* in acidic environments was described by Ahmed et al. [33] and was associated with phenotypic heterogeneity, but no mention of PHAs was included in that work. In addition, Moradali and Rhem [34] reported that there is scarce information about the effect of PHAs on bacterial persistence and dormancy, even in the context of highly studied topics such as pathogeny. In this work, the slight growth observed after inoculation at pH 5.5 could be related to the PHA accumulation heterogeneity in the starter cultures. The survival and replication of the cell subpopulation that had accumulated PHA may be responsible for the observed growth. It has been described that PHA depolymerization enhances the levels of the master regulator of stress response, RpoS [35]. This stationary phase Sigma factor has been related to protection against several stressors, including low and high pH [36,37]. In this case, it is possible that the accumulated PHA was depolymerized during the acidic pH shock, increasing the RpoS cell pool and leading to acidic pH resistance. A study performed in a recombinant *E. coli* strain showed that the capability to accumulate and depolymerize PHB enhanced bacterial survival to acidic pH, with both *phbC* and *phaZ1*, encoding PHB polymerase and depolymerase, respectively, being necessary for the highest survival [16]. Accordingly, our results regarding different mutants showed that the PHAB^-^ strain, which was unable to accumulate PHAs, was the most sensitive to pH stress. On the other hand, the pH sensitivity of the PHA^-^ mutant, which was able to produce only sclPHAs, resulted similar to the wild-type strain, indicating the relevance of sclPHA on the survival of this bacterium. Interestingly, it was described that *P. extremaustralis* sclPHA genes are located in a genomic island, probably derived from β-Proteobacteria [38], showing the importance of these horizontally transferred acquired genes also for pH adaptability.

We also observed that, at the end of the experiment, the extracellular pH of both, acidic and alkaline cultures was modified, converging in a pH around 8.6. The culture media alkalization was previously observed when ammonia was released due to peptones’ use as carbon and nitrogen sources [39], in this case from LB medium. Interestingly, cultures inoculated under alkaline conditions showed a decrease in media pH at the end of the assays. This pH decrease could be related to dynamic PHA synthesis and degradation [4, 6]. During PHA depolymerization two events occur: the monomers’ utilization as carbon and reducing power source and the extracellular release of both (R)-3-hydroxybutyric (3HB) and mcl(R)-3-hydroxyalkanoic (3HAmcl) acid [40,41]. In *P. putida* previous studies showed that alkaline pH increases the activity of the intracellular mclPHA depolymerase (PhaZ) leading to monomers liberation with the consequent decrease of the extracellular pH [41]. However, near neutral pH, the monomer liberation is less efficient [42]. In *P. extremaustralis*, when the initial pH was 9.0 and 9.5, the pH of both supernatants after 24 h dropped to 8.6, but with an initial pH of 10.0, the pH decreased to 9.0 at the end of the experiment. We also observed that pH changes were related to bacterial growth, since when cultures showed a scarce growth rate, extracellular pH remained similar to the initial one. Traditionally, PHA-producing bacteria have attracted biotechnological interest due to their potential applications in bioplastic production. Besides, it was widely demonstrated that PHAs have ecological importance due to their role in bacterial fitness conferring additional benefits for other environmental applications [3-5]. Thus, PHA accumulation has been useful in tolerating different stresses such as UV-radiation, heat, osmotic shock, desiccation, and oxidative agents in plant growth-promoting bacteria and also in coping with contaminant-derived stress in microorganisms used for bioremediation purposes [3, 4]. The bacterial activity during the bioremediation process can change the pH of hydrocarbon-contaminated soils and consequently influence soil parameters such as nutrient availability, solubility contaminant, and bioavailability required for microbial metabolism [43]. Optimum soil pH is important to regulate microbial biomass and enzyme activity necessary for hydrocarbon biodegradation [44]. It has been reported that it is important to adjust the soil pH in hydrocarbon-contaminated sites ranging from 5.0 to 8.0 for the success of bioremediation [43]. The same landscape can be found in agricultural soils. Excessive use of nitrogen fertilizer and acid precipitation prompt soil acidification whereas soil alkalization is mainly due to the input of alkaline oxide dust and ash [45,46]. The acidification and alkalization of soils is a worldwide problem that has been increased by human activities, thus affecting crop yield as well as microbial diversity [44]. Then, PHA accumulation can be a key factor supporting bacterial survival by alleviating pH-derived stress in these environments.

## Conclusions

Our analyses showed that pH values ranging from 6.0 to 9.5 supported growth and PHA accumulation in *P. extremaustralis* 14-3b. Furthermore, we observed that intracellular PHA content, particularly sclPHA, provided protection against both acidic and alkaline pH stress. This discovery highlights the potential to enhance stress responses through tailored growth conditions for PHA-producing bacteria. These findings open new avenues for biotechnological and environmental applications, particularly in improving the survival of microorganisms in acidic or alkaline environments.

## Acknowledgments

This work was supported by grants from Consejo Nacional de Investigaciones Científicas y Técnicas-CONICET-(grant number: PUE14120210300619CO and PIP 11220200101436CO), and Agencia I+D (grant numbers: PICT2019 N°02482 and PICT2018-02319) from Argentina. NIL and LJRI are career investigators from CONICET.

## Conflict of Interest

The authors declare that they have no known competing financial interests or personal relationships that could have appeared to influence the work reported in this paper.

## CRediT Author Statement

M.G. Brito: Methodology, Formal analysis, Investigation. N.I. López: Conceptualization, Methodology, Writing – original draft, review & editing, Resources, Funding acquisition. L.J. Raiger Iustman: Conceptualization, Methodology, Supervision, Writing – original draft, review & editing, Resources, Funding acquisition.

## Notes

### Competing Interest Statement

The authors have declared no competing interest.

